# Basal EFNA3 Facilitates Luminal-Driving Mammary Epithelial Migration and Cancer Metastasis by Promoting OXPHOS

**DOI:** 10.1101/2024.10.18.619009

**Authors:** Jing Chen, Rongze ma, Deyi Feng, Zhixuan Deng, Yunzhe Lu, Kun Xia, Ophir D. Klein, Pengfei Lu

## Abstract

Collective migration is essential for tissue development, repair, and cancer metastasis. However, the mechanisms by which different cell types coordinate to facilitate cohesive movement remain poorly understood. Here, we report distinct migratory behaviors in luminal and basal cells during collective migration of the mammary epithelium. We show that luminal cells are leaders of directional migration. However, basal cells also play a central role in promoting collective migration by producing EFNA3, which targets luminal cells and enhances oxidative phosphorylation and energy production. This promotion of epithelial migration and branching morphogenesis is confirmed in vivo. Importantly, EFNA3 is a prognostic marker for breast cancer metastasis, with high expression correlating with increased metastasis. Conversely, inhibition of EFNA3 significantly slows metastasis. These findings reveal a novel mechanism of epithelial migration that is critical for organogenesis and identify EFNA3 as a potential therapeutic target for cancer metastasis.

## INTRODUCTION

Collective cell migration, where cells coordinate their movements as a group, is increasingly recognized as essential for various biological processes, including organ formation, tissue regeneration, wound healing, and disease progression^1,2^. Recent studies have demonstrated that collectively migrating cancer cells not only metastasize more efficiently but also exhibit enhanced survival at distant sites than single cells^3,4^. However, much of our understanding of collective migration stems from research on mesenchymal systems, which employ mechanisms distinct from those used by epithelial tissues ^5^. Given that epithelial tissues are the origin of approximately 90% of cancers, elucidating the mechanisms by which epithelial cells migrate during development and metastasis is a critical area of research^6–8^.

In all systems of directional migration, whether individual or collective, two fundamental questions arise: how is directionality (front-rear polarity) established, and what is the driving force behind migration^9^? In collective migration, a third key question emerges: how do cells within the collective collaborate to achieve coordinated, cohesive movement? Mesenchymal and epithelial systems address these questions through distinct mechanisms. In mesenchymal systems, leader cells—a fixed population at the migration front—direct collective movement by extending actin-rich filopodia or lamellipodia into the extracellular matrix (ECM), driving the migration of both leader and follower cells ^10^. Follower cells, far from passive, actively interact with leader cells, functioning in concert with the entire collective as if part of a larger “supracell”^11,12^.

In contrast, studies based on in vitro models reveal that mammary epithelial migration follows a two-phase process. Initially, stationary epithelial cells proliferate asymmetrically and stratify at the migration front, establishing front-rear polarity and generating leader cells^5^. In the second phase, leader and follower cells coordinate to drive collective migration (Supplementary Fig. 1A) ^5,13^. Notably, leader cells in the mammary epithelium constitute a dynamic population, continually changing positions—a sharp contrast to the fixed leader populations observed in mesenchymal tissues^5^. Additionally, these epithelial leader cells do not extend matrix-invading protrusions, a hallmark of mesenchymal migration^14^.

Crucially, the in vitro migratory behavior of mammary epithelial cells closely mirrors the in vivo structure and dynamics of terminal end buds (TEBs)—the branching tips of mammary glands—at the level of cellular composition, organization, and molecular signatures^5^. The relevance of this in vitro model is further underscored by findings that genetic ablation of *Lgl1*, a key tissue polarity gene, impairs both in vitro epithelial migration and in vivo epithelial branching^15^. This highlights the critical role of collective migration in normal mammary gland development and the utility of the in vitro system for exploring epithelial migration mechanisms applicable in vivo.

Recent studies further indicate that mammary epithelial migration operates independently of epithelial-mesenchymal transition (EMT), and that *Snail1*, a transcription factor ascribed to be essential for EMT, is instead crucial for regulating the directionality of collective migration ^16^. A deeper understanding of epithelial migration mechanisms is therefore essential for advancing knowledge of vertebrate organ development and cancer metastasis. The mammary epithelium, as a model for vertebrate glandular epithelial tissues, comprises two distinct cell populations—basal and luminal cells—each with unique morphologies and molecular profiles^17–19^. Here, we hypothesize that these two epithelial populations cooperate during collective migration, both in the context of epithelial branching and cancer metastasis. To test this hypothesis, we analyzed the behavior of individual basal and luminal cells and investigated the mechanisms underlying their interactions during development and cancer progression.

## RESULTS

### Basal Cells Exhibit Lower Adhesion and Higher Speed than Luminal Cells During Collective Migration

To assess the behavior of individual basal and luminal cells, we employed fluorescence-assisted cell sorting (FACS) to isolate green fluorescently labelled basal and luminal cells from mammary gland epithelium and tracked them during migration. These green basal and luminal cells were derived from mammary epithelial organoids of female mice expressing the *R26R*^mTmG^ (red) alleles infected with adenovirus Cre, which, upon infection and Cre expression, started to express membrane-GFP (green). They were then combined with red fluorescent cells from the mTmG mammary gland at a 1:9 ratio and were tracked during mammary organoid epithelial migration (Fig. 1A).

**Figure 1:**
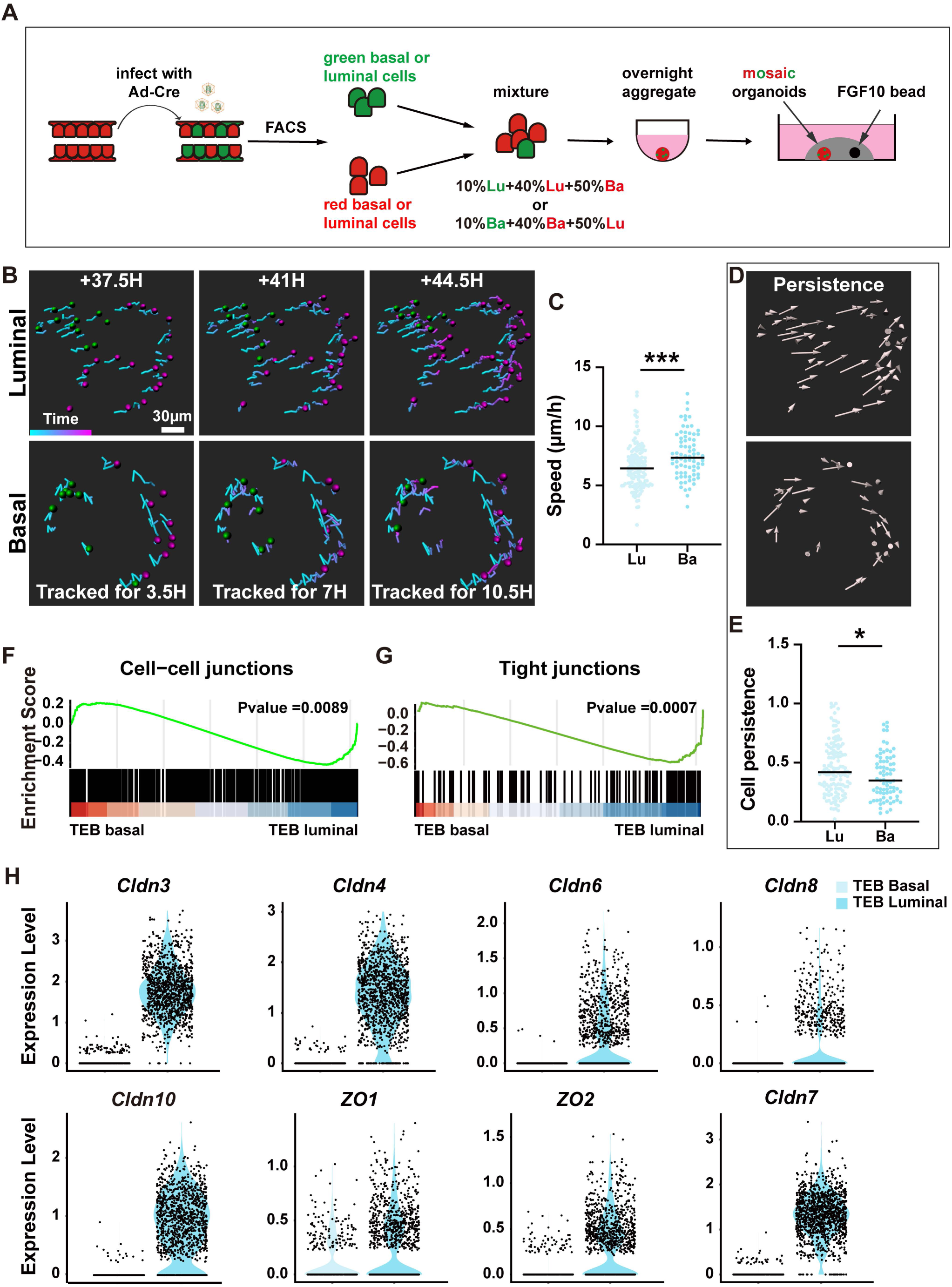
Basal Cells Exhibit Lower Adhesion and Higher Speed than Luminal Cells During Collective Migration. (**A**) Schematic diagram illustrating the experimental workflow for sample preparation and labeling of individual cells for time-lapse confocal imaging. Mammary organoids derived from *R26R*^mTmG^ mice were used as the source of red fluorescent luminal and basal cells. Green, fluorescent basal or luminal cells, used for tracking, were generated by infecting red *R26R*^mTmG^ mammary epithelial cells with adenovirus-Cre. Green fluorescently labeled basal or luminal cells were combined with red fluorescent cells from the *R26R*^mTmG^ mammary gland at a 1:9 ratio and tracked during mammary organoid epithelial migration. (**B**-**E**) Migration analyses, including cell tracks (**B**), mean speed (**C**), and persistence (**D**, **E**) of individual luminal (133 tracks from 4 organoids) and basal cells (76 tracks from 4 organoids). Persistence, defined as a cell’s tendency to maintain its direction of movement over time, was calculated from cell trajectories as displacement divided by total track length. Data are presented as mean ± SD. Statistical analysis was conducted using unpaired Student’s t test. *p < 0.05. Abbreviations: Ad, adenovirus; Ba, basal; Lu, luminal; TEB, terminal end buds. (**F**, **G**) GSEA analyses of the cell-cell junction (**F**) and the tight junction (**E**) pathways in basal cells and luminal cells within TEBs. Both pathways show statistically significant p values. (**H**) Expression of a panel of tight junction components in basal cells and luminal cells of TEBs, based on published scRNA-seq databases^20^.

Cell tracking from 37.5 to 44.5 hours revealed that basal cells migrate faster than luminal cells, with basal cells exhibiting a mean speed of ∼7 µm/h, while luminal cells moved at ∼4 µm/h (Fig. 1B, C; Movies 1, 2). However, despite their higher speed, basal cells demonstrated lower persistence, calculated as displacement/track length, compared to luminal cells (Fig. 1D, E).

We hypothesized that basal cells might have lower cell-cell adhesion, corresponding to their increased speed. Using published single cell RNA sequencing (scRNA-seq) data from TEBs of branching mammary epithelium ^20^, we performed Gene Set Enrichment Analysis (GSEA) and confirmed that basal cells exhibit lower expression of cell-cell junction components compared to luminal cells (Fig. 1F). Interestingly, despite the subtype-specific expression of adhesion components, the overall expression of adherens junction and desmosome components—except for a few—was not significantly different between the two cell types (Supplementary Fig. 1B-E). Notably, basal cells showed reduced expression of tight junction (TJ) components, most of which were luminal cell-specific (Fig. 1G, H).

In summary, basal cells move faster than luminal cells during migration, which correlates with reduced cell-cell adhesion, particularly tight junction components. Interestingly, luminal cells show higher persistence, suggesting a critical role in directional migration of the mammary epithelium.

### Frontal Luminal Cells Lead Mammary Epithelial Migration

Previous research indicated that leader cells at the front of migrating mammary epithelial organoids respond to the chemotactic signal FGF10 by preferentially extending intra-epithelial protrusions toward the signal ^5^. To identify the epithelial subtype giving rise to these leader cells, we analyzed the direction of cellular protrusions by assigning them to 45° bins relative to the axis of migration (Fig. 2A-D). Our analysis revealed that luminal cells at the front of the migrating epithelium exhibited a significant preference for FGF10 beads (Fig. 2A, C), while rear luminal cells (Fig. 2D) and basal cells (Fig. 2B, E, F) showed no such directional bias.

**Figure 2:**
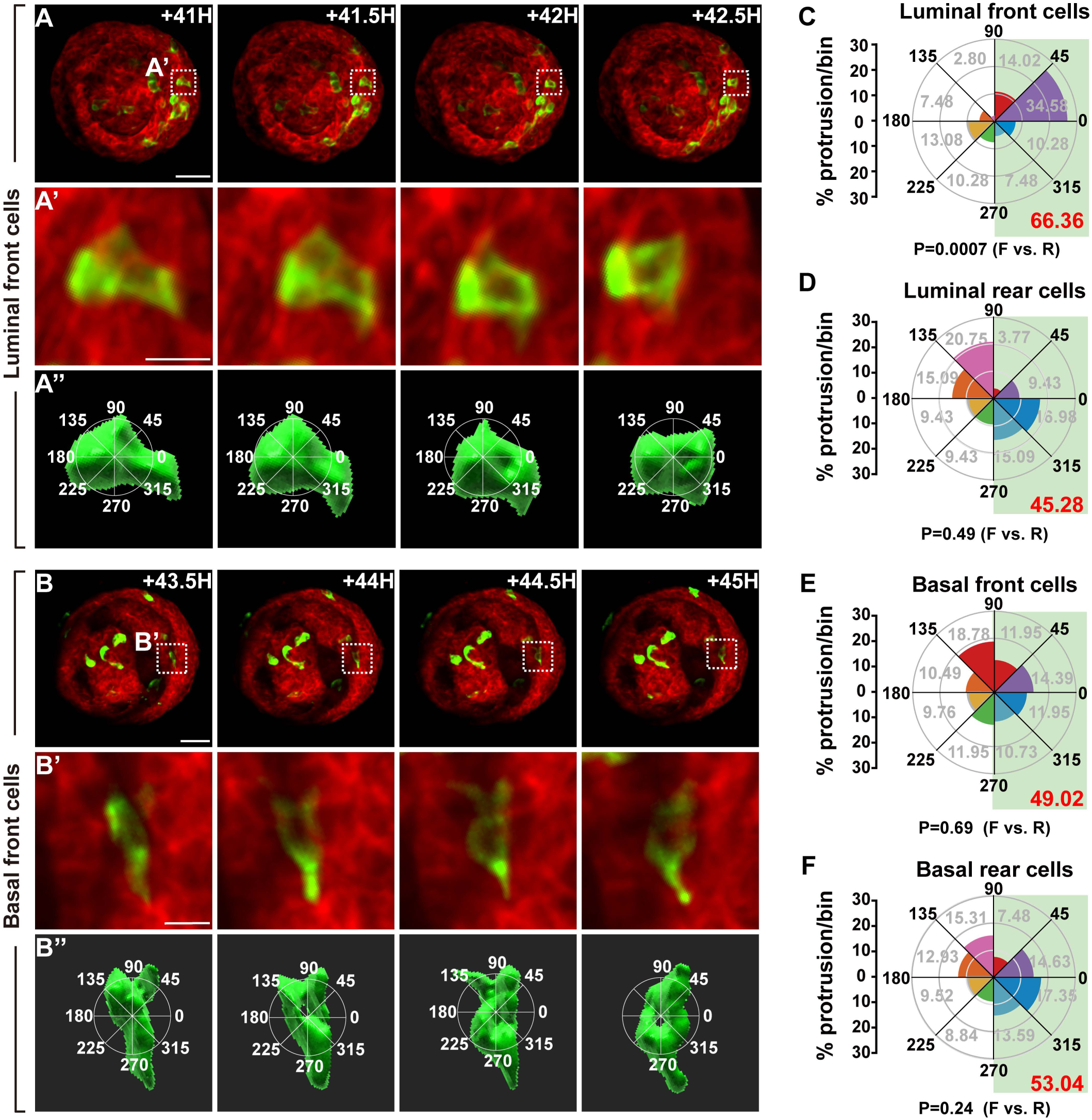
Frontal luminal Cells Lead Mammary Epithelial Migration. (**A**–**B’’**) 3D reconstructions of labeled front luminal (**A**–**A’’**) and basal (**B**–**B’’**) cells with protrusions, imaged during collective migration of mosaic organoids induced by FGF10 at the indicated time points. (**A’’**, **B’’**) Protrusions were assigned to 45° bins, with the 0–180° axis aligned along the migration direction. Scale bars, 20 µm. (**C**–**F**) Quantification of protrusions per bin from front luminal (**C**), rear luminal (**D**), front basal (**E**), and rear basal (**F**) cells of the mosaic organoid. To assess the total frequency of cell protrusions extending toward the front, protrusions observed between 90° and 270°, highlighted in the light green shade, were grouped as “front.” Total frequencies are indicated in the lower right corner of each panel. Chi-squared (χ²) tests were performed to determine if the difference in protrusion frequencies between the front (**F**) and rear (**R**) was statistically significant. The P value is significant in (**C**), but not in (**D**–**F**). For front luminal cells, a total of 3 organoids, including 107 protrusions, were analyzed (**C**). For rear luminal cells, 3 organoids with 53 protrusions were analyzed (**D**). For front basal cells, 4 organoids with 410 protrusions were analyzed (**E**). For rear basal cells, 4 organoids with 294 protrusions were analyzed (**F**). Abbreviations: F, front; R, rear.

These results suggested that luminal cells at the leading edge are responsible for driving directional migration, consistent with our earlier model proposing that luminal epithelium undergoes stratification and apicobasal polarity conversion to front-rear polarity, setting up migration directionality.

### The Focal Adhesion Pathway is Active in Luminal Cells

It is paradoxical that luminal cells lead mammary epithelial migration, as they are thought to be enveloped by basal cells and separated from the extracellular matrix (ECM), which provides the traction required for migration ^15,17,21^. We hypothesized that luminal cells interact with the ECM and possess active integrin signaling. Mining scRNA-seq data from the developing mammary epithelium ^20^, we found that the expression level of the focal adhesion pathway is indistinguishable between basal and luminal cells in the TEBs based on the GSEA (Supplementary Fig.2A). When individual genes of the pathway were examined, we found that they all, including *Itga2*, *Itgb1*, *Itgb3*, and *Fak*, were expressed in both basal and luminal cells (Fig. 3A; Supplementary Fig. 2B). qPCR analysis further validated these results in 8-week-old mammary glands (Fig. 3B; Supplementary Fig. 2C). Interestingly, while *Itgb1* expression was higher in basal cells, *Itga2*, *Itgb3*, and *Fak* were more highly expressed in luminal cells (Fig. 3B).

**Figure 3:**
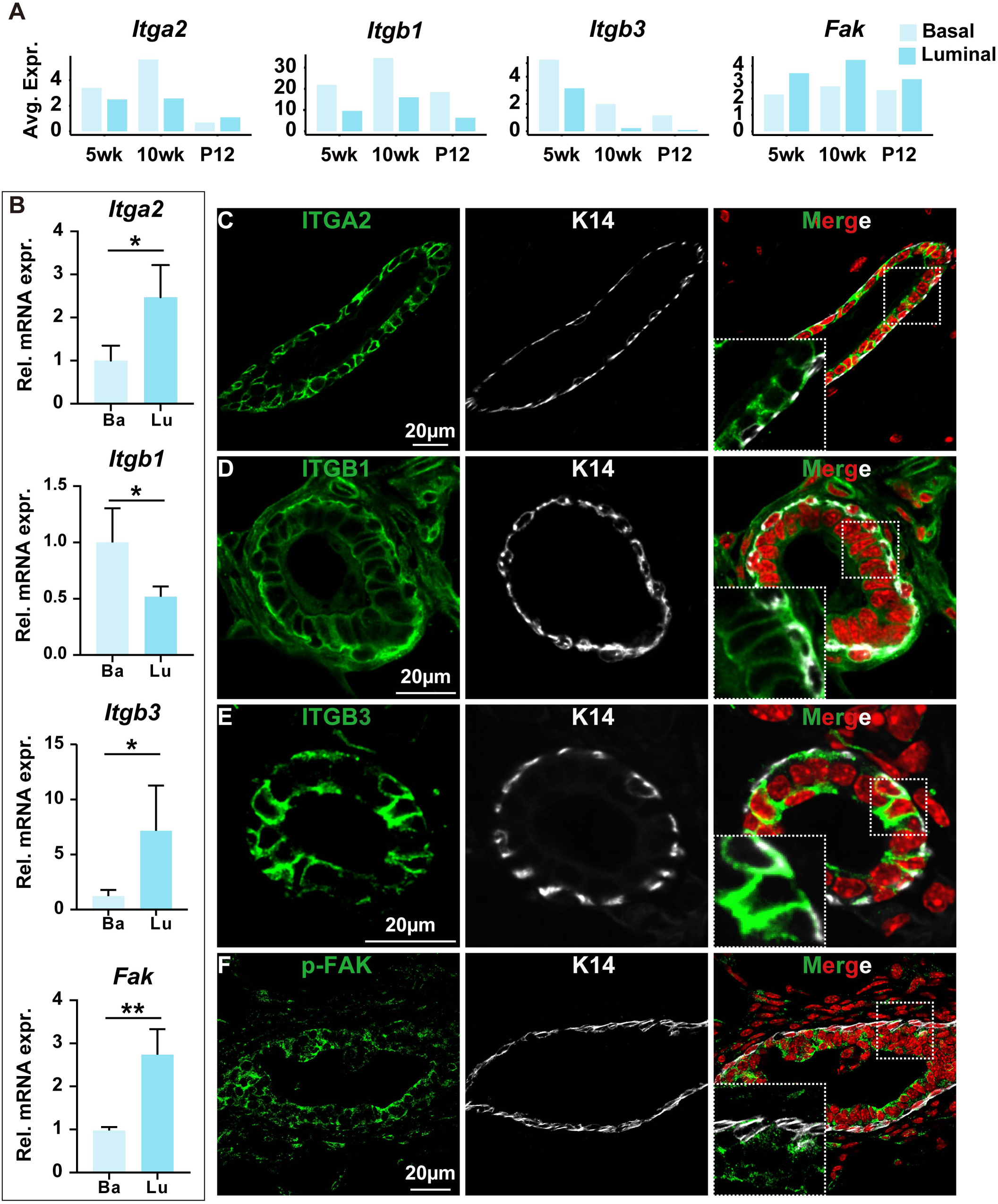
The Focal Adhesion Pathway Is Active in Luminal Cells. (**A-F**) mRNA expression (**A**, **B**) and protein expression (**C**-**F**) of selected members of the focal adhesion pathway in basal and luminal cells during mammary gland development at the indicated stages. (**A**) Results are based on published single-cell RNA sequencing data (GSE164017). (**B**) mRNA expression was detected by qPCR. (**C**-**F**) Protein expression and localization were assessed using immunofluorescence microscopy, employing antibodies specified in the Methods section. K14-GFP mice were used to visualize basal cells (white). Data are presented as mean ± SD. n.s. not significant, P ≥ 0.05; *P<0.05, ** P < 0.01. Abbreviations: wk, week; P, pregnancy day; rel, relative; expr, expression.

Immunofluorescent confocal microscopy confirmed protein expression in luminal cells, and phospho-FAK staining indicated that the integrin signaling pathway is active (Fig. 3C-F). These data indicated that luminal cells, despite being encapsulated by basal cells, can interact with the ECM and have active focal adhesion signaling.

### Basal Cells Promote Luminal Cell-Driven Collective Migration by Increasing Speed and Stabilizing Directionality

Given that luminal cells drive mammary collective migration, we hypothesized that mammary epithelial aggregates composed exclusively of luminal cells would migrate more efficiently than those containing both luminal and basal cells. To test this, we used flow cytometry to purify luminal and basal cells and generated mammary aggregates of three types: luminal-only (Lu), basal-only (Ba), and mixed luminal-basal aggregates (BaLu) (Fig. 4A). In migration assays, both luminal and basal aggregates exhibited migration capabilities (Fig. 4B, C). However, to our surprise, aggregates containing both cell types migrated more efficiently than either luminal- or basal-only aggregates, traveling greater distances and at a faster pace (Fig. 4D, E). These findings indicate that basal cells actively contribute to, rather than impede, luminal cell-driven collective migration.

**Figure 4:**
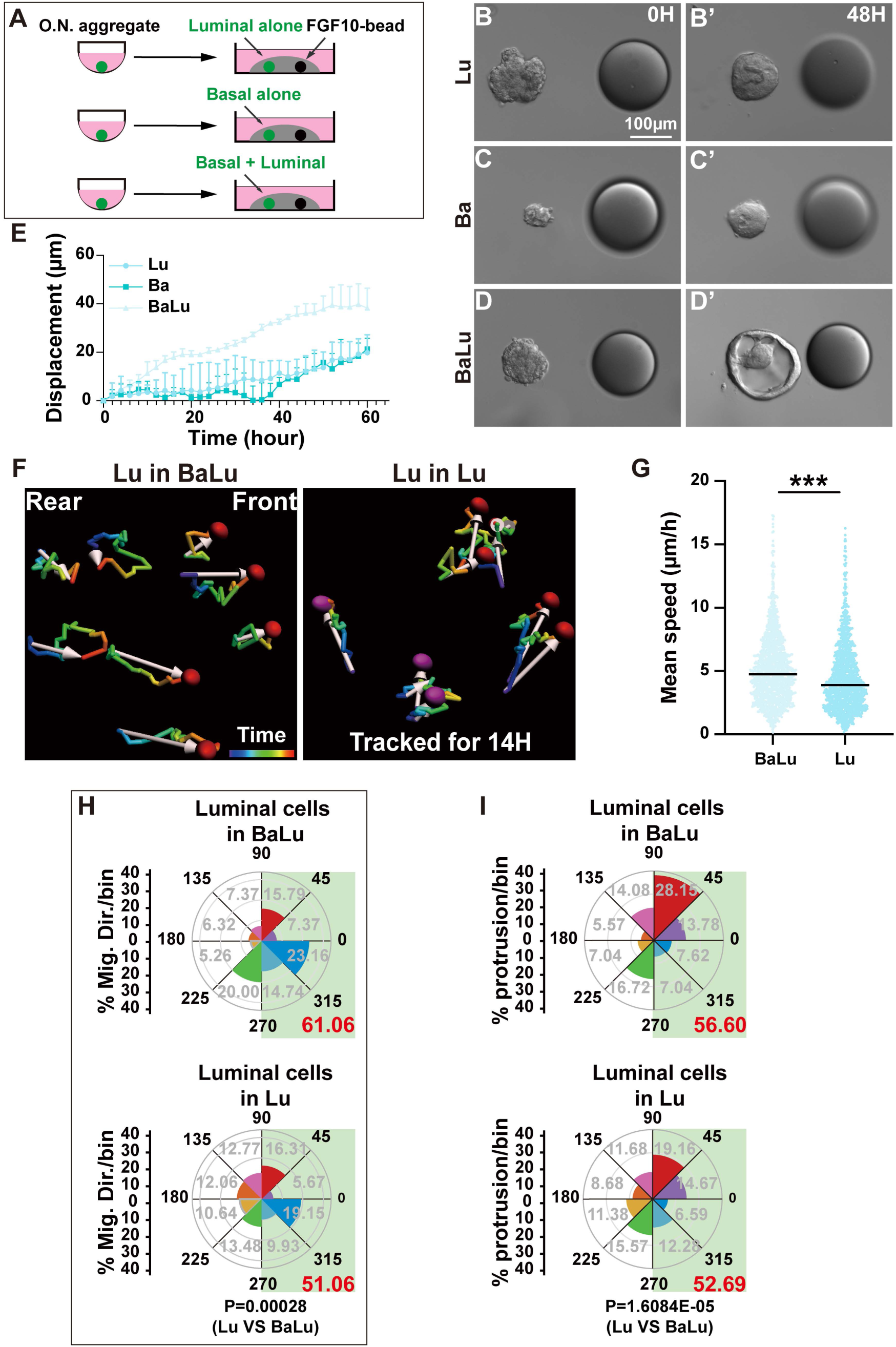
Basal Cells Promote Luminal Cell-Driven Collective Migration by Increasing Speed and Stabilizing Directionality. (**A**) Schematic diagram illustrating the experimental workflow for organoid preparation, including basal and luminal cell sorting, aggregation, and in vitro culture methods. Approximately 2000 cells were used to form each aggregate. (**B**-**D**) Still DIC images of epithelial aggregates composed of luminal cells alone (abbreviated as Lu, **B**, **B’**), basal cells alone (abbreviated as Ba, **C**, **C’**), or a mixture of basal and luminal cells (abbreviated as BaLu, **D**, **D’**) during migration. Scale bars: 100 μm. (E) Quantification of migration distance for luminal and basal cell aggregates. (**F**-**H**) Migration analyses, including cell tracks (**F**), mean speed (**G**), and migration direction (**H**) of individual luminal cells in either basal-luminal cell mixtures (BaLu; 95 tracks from 5 organoids) or luminal-only aggregates (Lu; 141 tracks from 5 organoids). (**F**) Cells were tracked for 14 hours. White arrows indicate the starting and ending positions of each tracked cell, with the arrow lengths representing the distance traveled over the time span. (**G**) Mean cell speeds were calculated by dividing the track length by time. (**H**) Migration direction per bin was quantified for luminal cells in BaLu mixtures or luminal-only aggregates. Total frequencies are indicated in the lower right corner of each panel. (I) Quantification of protrusions per bin for luminal cells in BaLu mixtures and luminal-only aggregates. A total of 4 BaLu organoids (764 protrusions) and 4 Lu organoids (583 protrusions) were analyzed. Total frequencies are indicated in the lower right corner of each panel. Chi-squared (χ²) tests were used to compare the frequency of luminal cells migrating toward the front or rear. Note that the P value is significant. Data were mean ± SD. *** P < 0.001. Abbreviations: O.N., overnight; Ba, basal; Lu, luminal; mig, migration; dir, direction; VS, versus.

To elucidate how basal cells influence luminal behavior, we labeled luminal cells and tracked their migration speed and morphology in the presence and absence of basal cells. We observed that luminal cells at the migration front moved faster in mixed aggregates compared to luminal-only aggregates (Fig. 4F, G). Furthermore, luminal cells displayed a greater preference for aligned, directional movement when basal cells were present (61.06% vs. 51.06%) (Fig. 4H). These results demonstrate that basal cells enhance both the speed and directional persistence of luminal cells during collective migration. Consistent with these observations, we found that, although frontal luminal cells preferentially extended protrusions toward FGF10 beads in both the presence and absence of basal cells, those in mixed aggregates did so at a higher frequency (56.6%) compared to luminal-only aggregates (52.69%) (Fig. 4I).

Together, these results indicated that basal cells facilitate luminal cell-driven collective migration by increasing both migration speed and directional persistence.

### *Efna3* Facilitates Epithelial Migration via the OXPHOS Pathway

To elucidate the molecular mechanisms by which basal cells promote organoid migration, we performed bulk RNA-seq on migrating aggregates composed of either luminal or basal cells alone, or a combination of both cell types. Transcriptomic analysis identified 2,462 genes specific to basal cells in the Ba group, while 2,135 genes were upregulated in the BaLu aggregates containing both basal and luminal cells. By subtracting the 1,323 genes that were also increased in the Ba group from the upregulated genes in the BaLu group, we identified 812 genes that were primarily upregulated in luminal cells due to stimulation by basal cells (Fig. 5A).

**Figure 5:**
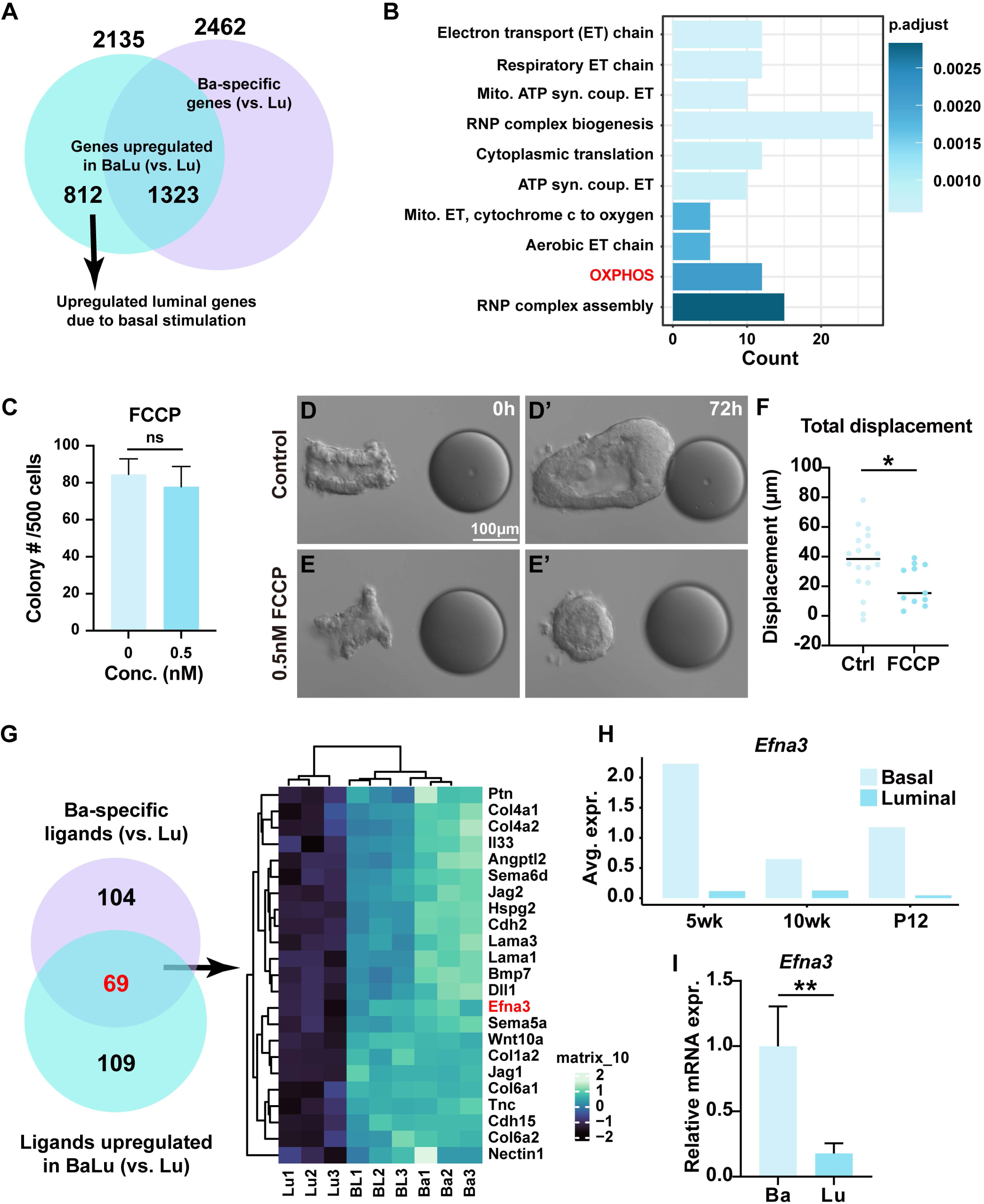
*Efna3* Facilitates Epithelial Migration via the OXPHOS Pathway. (A) Venn diagram illustrating the bioinformatics analysis that led to the identification of genes upregulated in luminal cells following stimulation by basal cells. Bulk RNA-seq was used to compare gene expression between the BaLu group and the Lu group to isolate genes upregulated by the former. To exclude genes specifically expressed by basal cells, a comparison between Ba and Lu was performed, and genes highly expressed in the Ba group were subtracted from those upregulated in the BaLu group. (B) Gene Ontology (GO) analysis showing the main pathway alterations in luminal cells, either in the absence (control luminal-alone aggregates) or presence (experimental BaLu aggregates) of basal cells. Notably, pathways related to energy production, including electron transport chain and OXPHOS are strongly represented. (C) Effect of the OXPHOS inhibitor FCCP on luminal cell colony formation. (**D**-**F**) Impact of FCCP on FGF10-induced epithelial migration. FCCP was added to the medium at the indicated concentrations (**D**-**E’**). The effect of FCCP at 0 nM (n=18) and 0.5 nM (n=10) was quantified (**F**). (G) Top 25 basal-specific ligands potentially targeting luminal cells via their cognate receptors expressed in luminal cells. To derive this list, we first identified 109 significantly expressed ligands in the BL vs. Lu comparison. Similarly, the Lu vs. Ba comparison revealed 104 Ba-specific ligands. Intersecting these two sets yielded 69 ligands specifically expressed in Ba, which may influence Lu during the migration process. (H) mRNA expression of *Efna3* in basal and luminal cells at different stages of mouse mammary gland development, based on scRNA-seq dataset analysis(GSE164017). (I) Relative expression of *Efna3* in sorted basal and luminal cells. Data are presented as mean ± SD. n.s. = not significant, P ≥ 0.05; *P < 0.05, **P < 0.01. Abbreviations: Mito, mitochondria; syn, synthesis; coup, coupled; conc, concentration; wk, week; P, pregnancy day.

Gene Ontology (GO) analysis of these upregulated luminal genes highlighted significant enrichment in key pathways associated with cellular respiration and energy production, particularly oxidative phosphorylation (OXPHOS) and the electron transport (ET) chain (Fig. 5B). This analysis revealed enhanced mitochondrial ATP synthesis coupled with electron transport, with a specific focus on the aerobic electron transport chain, particularly from cytochrome c to oxygen. Concurrently, we observed an increase in ribonucleoprotein (RNP) complex biogenesis and cytoplasmic translation, suggesting a coordinated interaction between energy metabolism and protein synthesis. This pattern indicates an elevated demand for ATP, likely driven by increased cellular activity during epithelial collective migration, along with enhanced protein production to support these metabolic processes.

To determine whether OXPHOS is essential for epithelial migration, we first identified a tolerable concentration of FCCP (carbonyl cyanide 4-fluoromethoxyphenylhydrazone), a mitochondrial uncoupler that disrupts the proton gradient across the inner mitochondrial membrane. At a concentration of 0.5 μM, FCCP did not affect colony formation (Fig. 5C). Subsequently, we added either a blank control or FCCP (0.5 μM) to the medium and monitored mammary organoid epithelial migration. FCCP treatment inhibited mammary epithelial migration by over 60%, confirming the critical role of OXPHOS in this process (Fig. 5D-F).

To determine whether OXPHOS is essential for epithelial migration, we first identified a tolerable concentration of FCCP (carbonyl cyanide 4-fluoromethoxyphenylhydrazone), a mitochondrial uncoupler that disrupts the proton gradient across the inner mitochondrial membrane. At a concentration of 0.5 nM, FCCP did not affect colony formation in luminal cells (Fig. 5C). Subsequently, we added either a blank control or FCCP (0.5 nM) to the medium and monitored mammary organoid epithelial migration. FCCP treatment inhibited mammary epithelial migration by over 60%, confirming the critical role of OXPHOS in this process (Fig. 5D-F).

Next, we focused on the top 25 basal-specific ligands likely mediating basal-luminal paracrine signaling (Fig. 5G). These include genes encoding cell adhesion molecules such as *Nectin1* and *Cdh15*; ECM components like *Col6a2*, *Col1a2*, and *Hspg2*, and growth factors such as *Jag1*, *Wnt10a*, and *Bmp7*, the latter of which we previously identified as crucial basal factors promoting luminal expansion^22^. We decided to focus on *Efna3*, encoding a member of the Ephrin family with limited functional understanding. Given the Ephrin family’s role in bi-directional signaling, we hypothesized that EFNA3 might be involved in basal-to-luminal signaling during epithelial migration.

Using the published scRNA-seq databases mentioned above, we found that *Efna3* is enriched in basal cells during mammary gland development, especially from around puberty onset at five weeks of age when it is almost exclusively expressed by basal cells (Fig. 5H). By contrast, we found multiple Ephrin receptor genes, including *Epha1*, *Epha2*, and *Ephb6*, were expressed in luminal cells during mammary development (Supplementary Fig. 3A). We performed qPCR and confirmed that *Efna3* is highly enriched in basal cells, while its candidate receptors are expressed in luminal cells of the eight-week-old mammary gland during which active epithelial branching and expansion are ongoing (Fig. 5I; Supplementary Fig. 3B).

Together, these studies identified EFNA3, a member of the Ephrin family, as a candidate basal factor that promotes luminal collective migration.

### Basal EFNA3 Promotes Luminal Migration In Vitro and Epithelial Branching In Vivo by Enhancing OXPHOS

To investigate the function of *Efna3*, we knocked down its expression in basal cells using lentiviral shRNA, achieving a 75% reduction (Fig. 6A). Co-culture of these *Efna3*-deficient basal cells with wild-type luminal cells resulted in a significant impairment of collective migration (Fig. 6B-D), indicating that *Efna3* in basal cells is crucial for luminal cell migration. Next, we overexpressed *Efna3* in basal cells, which led to a 45-fold increase in mRNA levels (Fig. 6E). This overexpression markedly enhanced the collective migration of mammary aggregates (Fig. 6F-G).

**Figure 6:**
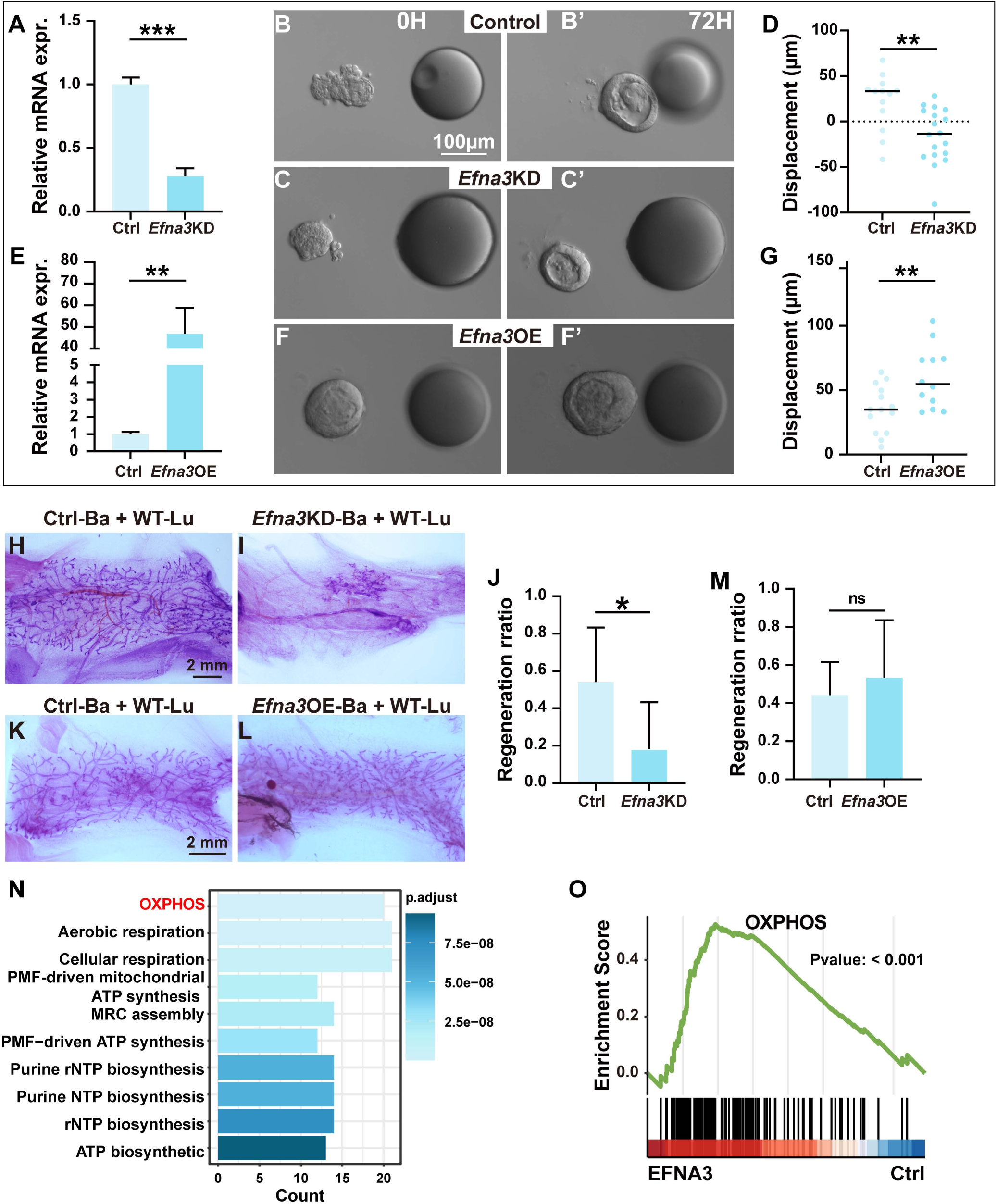
Basal EFNA3 Promotes Luminal Migration In Vitro and Epithelial Branching In Vivo by Enhancing OXPHOS. (**A**-**G**) Effect of *Efna3* loss- and gain-of-function on mammary epithelial migration, assessed using shRNA-based knockdown (KD) and an overexpression (OE) construct, respectively. (**A**, **E**) Relative *Efna3* mRNA expression in primary basal cells transfected with either a scrambled control or *Efna3*-KD lentiviral construct (**A**) or a control and *Efna3*-overexpression construct (**E**). (**B**-**G**) DIC still images from a time-lapse series of epithelial migration in representative control groups (**B**, **B’**), expressing scrambled shRNA or an empty vector as controls for *Efna3*KD or *Efna3*OE, respectively), the *Efna3*KD group (**C**, **C’**), and the *Efna3*OE group (**F**, **F’**). Quantification of epithelial displacement in the *Efna3*KD (**D**) and *Efna3*OE (**G**) experiments is shown. (**H**-**M**) Effect of *Efna3* loss- and gain-of-function in basal cells on mammary gland regeneration using the cleared fat-pad transplantation model. The epithelial network was visualized by Carmine staining in recombinant mammary glands derived from wildtype luminal cells mixed with basal cells transfected with either a control shRNA construct (**H**) or the *Efna3*KD construct (**I**) in the loss-of-function experiment, or with basal cells transfected with a control overexpression construct (**K**) or the *Efna3*OE construct (**L**) in the gain-of-function experiment. Quantification of regeneration in the *Efna3*KD (**J**) and *Efna3*OE (**M**) experiments was based on the regeneration ratio, defined as the ratio of the area covered by epithelial growth over the total area of the cleared fat pad. (**N**, **O**) KEGG (**N**) and GSEA (**O**) analyses of the upregulated genes as a result of EFNA3 protein stimulation on luminal cells after 24 hours of treatment. Data are presented as mean ± SD. n.s. indicates not significant (P ≥ 0.05); *P < 0.05; **P < 0.01; ***P < 0.001. Abbreviations: Ctrl, control; PMF, proton motive force; MRC, mitochondrial respiratory chain.

To assess the in vivo role of *Efna3*, we transplanted mammary epithelial cells (MECs) consisting of wild-type luminal cells and basal cells transfected with either control or *Efna3* shRNA into the cleared fat pads of three-week-old nude mice. Knockdown of *Efna3* in basal cells reduced mammary gland regeneration by 80%, underscoring the critical role of *Efna3* in epithelial branching (Fig. 6H-J). However, overexpression of *Efna3* in basal cells did not enhance mammary gland regeneration in vivo (Fig. 6K-M), suggesting that *Efna3* alone is insufficient to drive epithelial branching.

We then explored the mechanism by which *Efna3* functions. Consistent with our gain-of-function results, the addition of recombinant EFNA3 protein to culture media promoted luminal cell migration (Supplementary Fig. 4A-C). To further investigate, we performed bulk RNA-seq on luminal cells with and without EFNA3 treatment Gene ontology analysis of the upregulated genes in EFNA3-treated luminal cells revealed strong enrichment of pathways related to OXPHOS, aerobic respiration, cellular respiration, proton motive force (PMF)-driven ATP synthesis, and rNTP biosynthesis (Fig. 6N). This suggests a coordinated increase in energy production and nucleotide synthesis.

These findings indicate that EFNA3 stimulates heightened OXPHOS and mitochondrial activity, reflecting an active metabolic and proliferative state during epithelial migration. GSEA analysis further confirmed increased OXPHOS activity (Fig. 6O). In contrast, GO analysis of the downregulated genes in EFNA3-treated luminal cells highlighted pathways related to glycerolipid metabolism, female gonad development, and calcium signaling, though the relevance of these changes remains unclear (Supplementary Fig. 4D).

We further analyzed a panel of key genes involved in the OXPHOS pathway and confirmed their upregulation in EFNA3-treated cells (Supplementary Fig. 4E-G). Together, these results demonstrate that EFNA3 is essential for mammary epithelial migration in vitro and for branching morphogenesis in vivo, primarily by enhancing OXPHOS-related metabolism.

### EFNA3 as a Prognostic and Therapeutic Target to Inhibit Breast Cancer Metastasis

Given EFNA3’s role in epithelial migration, we predicted that it may be upregulated during cancer progression to promote metastasis. Indeed, analysis of the TCGA database revealed significantly elevated EFNA3 mRNA levels in breast cancer tissues (Fig. 7A). Using published scRNA-seq data on breast cancer subtypes ^23^, we found that EFNA3 is especially upregulated in the triple-negative breast cancer (TNBC) subtype (Fig. 7B). Moreover, high EFNA3 expression was associated with a significantly reduced probability of distant metastasis-free survival (DMFS, Fig. 7C). In parallel, OXPHOS levels were elevated in breast cancer tissues relative to normal tissue, as shown by GSEA and individual gene analysis in the OXPHOS pathway (Fig. 7D; Supplementary Fig. 5A). This upregulation positively correlated with increased EFNA3 expression (Supplementary Fig. 5B). Collectively, these correlative data support the hypothesis that EFNA3 promotes metastasis associated with OXPHOS in breast cancer patients.

**Figure 7:**
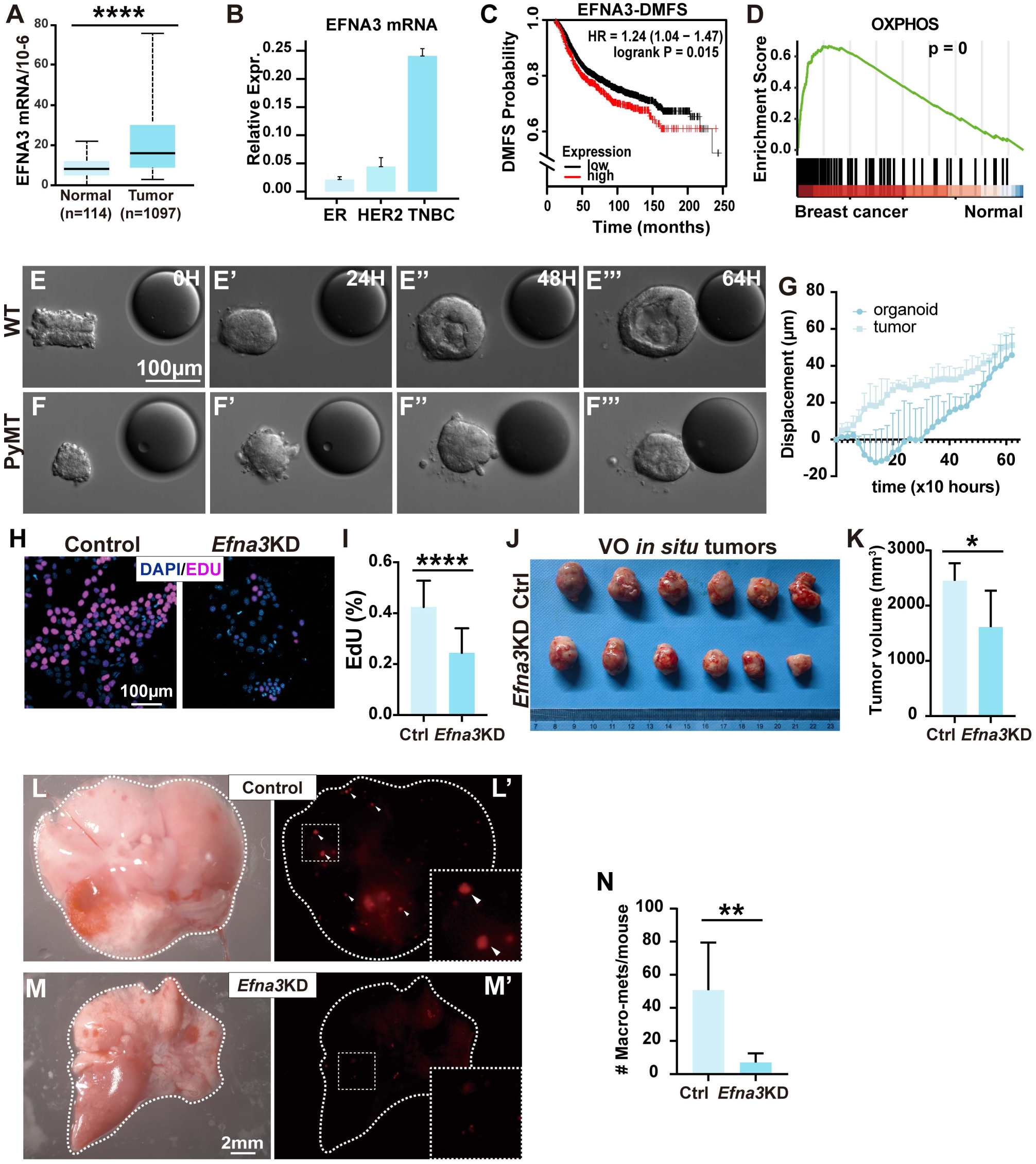
EFNA3 as a Prognostic and Therapeutic Target to Inhibit Breast Cancer Metastasis. (**A**, **B**) EFNA3 mRNA expression in human breast cancer overall based on TCGA database analysis (**A**) and across different subtypes based on scRNAseq data from PRJNA762594 (B). (C) Kaplan-Meier distant metastasis-free survival (DMFS) curve, showing that elevated EFNA3 expression correlates with reduced DMFS probability in patients. (D) GSEA analysis of the OXPHOS pathway in human breast cancer samples compared to normal samples. Data are sourced from TCGA. (**E**-**G**) Migration assays of organoids derived from wildtype (**E**-**E’’’**) and PyMT mammary epithelium (**F**-**F’’’**). Displacement of the organoids was quantified over time (hours) (**G**). Note lumen is absent from the tumor organoids, consistent with a loss of tissue polarity in the tumor tissue. Three wildtype organoids and four PyMT organoids were used for this assay. Scale bars: 100 μm. (**H**, **I**) Cell proliferation analysis of VO-PyMT cells transfected with a lentiviral construct expressing control scrambled shRNA or *Efna3* shRNA, measured by EdU incorporation (**H**). (**I**) Quantification of the percentage of EdU incorporation in control and *Efna3* knockdown cells. More than 500 cells were counted per sample. Scale bars: 100 µm. (**J**, **K**) Tumor growth in VO-PyMT cells 4 weeks post-surgery in nude mice, comparing control and *Efna3* knockdown conditions (**J**). Tumor volume is quantified in (**K**). (**L**-**N**) Lung metastasis from VO-PyMT cells transfected with lentivirus expressing control (**L**, **L’**) or *Efna3*KD shRNA (**M**, **M’**) observed in wholemount lung tissue under brightfield microscopy (**L**, **M**) and fluorescent stereoscope (**L’**, **M’**). (**N**) Quantification of lung metastasis from VO- PyMT cells transplanted into the cleared fat pad 4 weeks post-surgery. Data are presented as mean ± SD. *P < 0.05; **P < 0.01; ****P < 0.0001. Abbreviations: DMFS, distant metastasis-free survival; WT, wildtype; mets, metastases.

To experimentally test this hypothesis, we harvested mammary organoids from both wildtype and PyMT female mice and subjected them to an in vitro migration assay. PyMT organoids began migrating immediately at the onset of the assay, without the ∼30-hour lag phase observed in wildtype organoids and reached the FGF10 bead position earlier than the wildtype organoids (Fig. 7E-G), suggesting enhanced epithelial invasion in tumor tissue.

Next, we investigated the effects of downregulating *Efna3* during cancer progression. *Efna3* knockdown (KD) reduced the proliferation of mouse breast cancer cells by approximately 49%, as measured by the EdU incorporation assay (Fig. 7H, I). In an in vivo transplantation assay, PyMT cells with reduced *Efna3* expression exhibited a ∼60% reduction in tumor growth, both in volume and weight (Supplementary Fig. 5C-E). Since metastasis was not observed in either control or experimental host mice, we transitioned to the VO-PyMT line, a fluorescently and bioluminescently labeled cancer cell line derived from the PyMT model ^24^, for the metastasis assay.

VO-PyMT cells transfected with either control or *Efna3* shRNA lentivirus were transplanted into the cleared fat pads of six-week-old nude mice, and tumor growth and metastasis were examined four-weeks post-surgery. Compared to controls, VO-PyMT cells with *Efna3* knockdown exhibited an approximate 30% reduction in both tumor volume and weight (Fig. 7J, K; Supplementary Fig. 5F). At this stage, lung metastasis was readily detectable in mice transplanted with control VO-PyMT cells, with an average of 50 macro-metastases per mouse (Fig. 7L, L’, N). In contrast, mice transplanted with *Efna3* knockdown VO cells exhibited a significant reduction in macro-metastases, averaging only 5 per mouse—a 90% decrease (Fig. 7M, M’, N). Interestingly, overexpression of *Efna3* in VO-PyMT cells did not result in increased primary tumor growth (Supplementary Fig. 5G-I) or lung metastasis (Supplementary Fig. 5J-L), suggesting that *Efna3* overexpression alone is insufficient to drive cancer progression in vivo.

In summary, our data indicated that EFNA3 is essential for metastasis, served as a prognostic marker, and suggested that its inhibition may offer a therapeutic strategy to target breast cancer metastasis.

## DISCUSSION

The mechanisms underlying vertebrate epithelial migration remain poorly understood, particularly in terms of cellular interactions during the process. In this study, we identified differential movement patterns between luminal and basal cells during collective mammary epithelial migration. Notably, basal cells are essential in orchestrating this process, secreting EFNA3, which acts on luminal cells to promote OXPHOS and enhance energy production. This interaction drives epithelial migration and branching morphogenesis, a role that we validated through in vivo experiments (Fig. 8A). Additionally, *Efna3* emerged as a key prognostic marker in breast cancer, where its elevated expression is associated with increased metastasis. Inhibition of EFNA3, conversely, was shown to impede metastatic progression (Fig. 8A’). These results reveal an essential mechanism of epithelial migration that is critical for organ development and highlight *Efna3* as a promising therapeutic target for preventing cancer metastasis.

**Figure 8:**
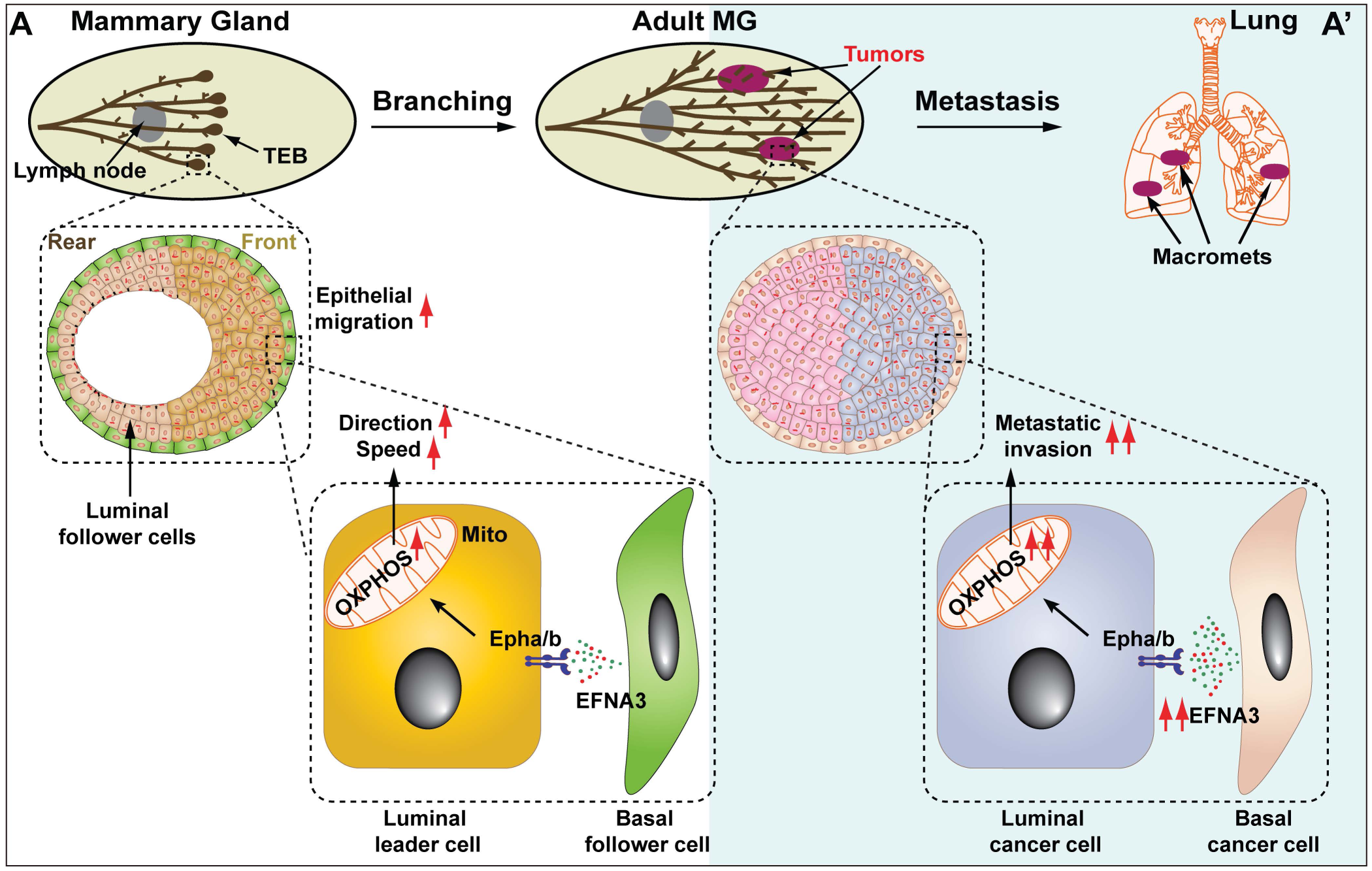
Model of Basal EFNA3 Function in Development and Caner Metastasis. (**A**, **A’**) Schematic diagrams depicting the role of EFNA3 in epithelial migration during development (**A**) and metastasis (**A’**). Previous findings demonstrated that epithelial migration within the terminal end buds (TEBs) is crucial for branching morphogenesis in vivo^15^. (**A**) In this study, we reveal that basal cells, along with rear luminal cells (as previously reported), act as follower cells, while frontal luminal cells serve as leaders in directional migration. Basal cells facilitate luminal-driven migration by secreting EFNA3, which binds to and activates receptors EPHA/B on luminal cells. This activation enhances OXPHOS activity, which is critical for the speed and directional stability of epithelial movement during organogenesis. (**A’**) In cancer progression, EFNA3 is upregulated, resulting in hyperactivation of the OXPHOS pathway in luminal cancer cells. This promotes increased epithelial invasion and metastasis to distant organs, such as the lungs. Our data suggest that EFNA3 is a promising therapeutic target for treating breast cancer metastasis. Abbreviations: MG, mammary gland; macro-mets, macrometastases; mito, mitochondrion.

### Luminal Cells as Leaders in Epithelial Migration

Several lines of evidence support the conclusion that luminal cells act as leaders in mammary epithelial migration. Specifically, the leading luminal cells exhibit high persistence, enabling them to maintain directional migration despite the dynamic nature of the epithelial front. In contrast, while basal cells move faster individually, they show less persistence. Consistent with these findings, luminal cells at the leading edge preferentially extend lamellipodia toward FGF10 compared to luminal cells further back or basal cells, regardless of their position. The preferential extension of cell protrusions in the direction of movement is a hallmark of leader cells during collective migration ^5^. Together, these data suggest that luminal cells have evolved specialized mechanisms to coordinate collective migration, ensuring that the epithelial sheet remains cohesive and directionally persistent during tissue remodeling and organogenesis.

Despite being enveloped by basal cells, luminal cells interact with the ECM via integrin signaling, further reinforcing their active role in migration. Our analysis revealed that integrin components, particularly ITGB1, ITGA2, and FAK, are expressed in luminal cells. Immunostaining confirmed active focal adhesion signaling, indicated by the presence of phospho-FAK, the active form of FAK in these cells. These findings suggest that luminal cells directly sense and respond to ECM cues during migration.

Our data showing dynamic movement of individual basal and luminal cells are consistent with the previous study illustrating frontal and rear cell behavior ^5^. This is in contrast to some mesenchymal systems where cells a in a fixed position and form a “supracell” to achieve cohesive movement ^11,12^, suggesting that the fundamental difference in how epithelium and mesenchyme collectively migrate. Moreover, previous study show that basal cells are at a leading position in both an in vitro model and in vivo tumor sample ^25^, our data suggest that this may be due to basal cells, having lower adhesion than luminal cells, thus occupying the leading position as a correlation, rather than evidence of being leader cells.

These data align with our previous reports showing that mammary epithelial migration occurs in two distinct stages. Initially, during the non-migrating phase, the front luminal epithelium undergoes proliferation-driven stratification and transitions from apicobasal to front-rear polarity, generating leader cells^5^. Although luminal cells lead collective migration in the mammary epithelium, with basal cells acting as followers, our data also show that basal cells can undergo directional migration in the absence of luminal cells. This suggests that basal cells can take on leader cell functions when luminal cells are absent.

### Basal Cells Promote Luminal Migration During Epithelial Branching and Lung Metastasis

While luminal cells lead migration, our data reveal that basal cells are crucial for facilitating luminal-driven collective migration. Basal cells increase the migration speed and directional persistence of luminal cells despite their own lower persistence. We demonstrated that basal cells produce EFNA3, which promotes OXPHOS in luminal cells—an essential energy source for migration. Transcriptomic analysis of migrating mammary organoids revealed significant upregulation of OXPHOS-related genes in luminal cells in the presence of basal cells, an effect abolished by *Efna3* knockdown. Moreover, treatment with FCCP, a mitochondrial uncoupler, inhibited mammary epithelial migration, confirming the central role of OXPHOS in this process.

These findings indicate a reciprocal relationship in which basal cells regulate the environment necessary for luminal cells to lead migration effectively. This is consistent with previous reports demonstrating that luminal cells inhibit basal stem cell differentiation ^26^ and our recent studies showing a positive feedback loop between basal and luminal epithelium, mediated by BMP7-INHBa signaling^22^. Together, these data underscore the importance of basal-luminal interactions in organ development and cancer progression.

The role of *Efna3* extends beyond development to cancer progression, where high *Efna3* expression correlates with breast cancer metastasis, particularly in aggressive subtypes such as the TNBC subtype. Our in vivo experiments demonstrated that *Efna3* knockdown in cancer cells significantly reduced tumor growth and metastasis. The reduction in lung metastasis observed in *Efna3*-knockdown animals compared to controls highlights the therapeutic potential of targeting *Efna3* in metastatic cancer. EFNA3’s ability to enhance OXPHOS in cancer cells mirrors its role in development, underscoring the similarities between developmental and cancerous epithelial migration. These findings align with studies identifying metabolic reprogramming as a key feature of metastatic cells, particularly their reliance on OXPHOS for energy production^27–29^.

Our study uncovers a complex interplay between luminal and basal cells during mammary epithelial migration, with luminal cells leading the process and basal cells providing essential support. The identification of EFNA3 as a basal-cell-derived factor that promotes OXPHOS in luminal cells adds a new dimension to our understanding of epithelial migration and cancer metastasis. EFNA3’s dual role in development and cancer progression makes it a promising therapeutic target for inhibiting breast cancer metastasis. Further research into *Efna3* and downstream signaling pathways may open new avenues for targeted interventions in both developmental disorders and metastatic cancers.

## MATERIALS AND METHODS

### Mouse Strains

Mice carrying the *R26R*^mT/mg.^ reporter allele (JAX Mice, #007576) ^24^, MMTV-PyMT (JAX Mice, #022974), K14-GFP transgenic mice (Cage Biosciences), wild-type C57BL/6JNifdc, and BALB/c Nude mice (GemPharmatechCo.,Ltd) were used. All mice were housed and maintained in compliance with the Institutional Animal Care and Use Committee (IACUC) guidelines at the University of South China.

### Mammary Organoid and Primary Epithelial Cell Isolation

Mammary organoids and primary epithelial cells were isolated using established protocols ^30^. The 2nd, 3rd, and 4th mammary glands were dissected, minced, and digested in 10 mL of digestion buffer (DMEM/F12 supplemented with 2 mg/mL collagenase [Sigma, C5138], 2 mg/mL trypsin [Gibco, 27250018], 5 µg/mL insulin [Yeasen, 40107ES25], 5% fetal bovine serum [Gibco, 10099141], and 50 µg/mL gentamicin) at 37°C with shaking for 25 minutes. The cell suspension was centrifuged at 560g for 10 minutes, followed by differential centrifugation at 450g to purify mammary organoids. Organoids were further dissociated using Trypsin-EDTA (TE) for 12 minutes to yield single cells.

### Epithelial Migration Assays

Migration assays for mammary gland and PyMT tumor organoids were conducted as previously described^30^ Primary mammary epithelial cells or virus-infected cells were diluted to a concentration of 20,000 cells/mL in mammary stem cell culture medium. A total of 100 µL of this suspension was plated in each well of a 96-well low-attachment plate and incubated overnight to form cell aggregates. Heparin sulfate beads (100-150 µm in diameter) were pre-soaked in 10 µL of 100 µg/mL FGF10 (GenScript, Z03155). After Matrigel (80%) (Corning, #354230) solidification on the culture dish, beads and mammary organoids were positioned ∼100 µm apart using a tungsten needle, followed by the addition of 1 mL basal medium (DMEM/F12 supplemented with 1% P/S, 1% L-Ala-Gln (Beyotime, C0211), and 1% ITS (Thermo, 41400045)). Live-cell imaging was performed to track migration dynamics. For the experiment involving EFNA3 protein (R&D, 7395-EA-050) stimulation of luminal aggregates, it is necessary to additionally add 1 µg/ml of EFNA3 protein to the basic medium.

### Cell Culture Conditions and Proliferation Assays

HEK293T cells(Pricella Life Science & Technology Co. Ltd., CL-0005)were cultured in DMEM (Gibco, # C12430500BT) supplemented with 10% FBS (Lonsera, S711-001S), 1% sodium pyruvate, 1% non-essential amino acids, 1% L-Ala-Gln, and 1% P/S. VO-PyMT cells were maintained in DMEM/F12 with 10% FBS, 1% P/S, 1 µg/mL hydrocortisone, 10 µg/mL insulin, and 1% L-Ala-Gln. Basal and luminal cells were cultured in DMEM/F12 supplemented with 5% FBS, 10 ng/mL EGF (Novoprotein, C029), 20 ng/mL FGF2 (GenScript, Z03116), 4 µg/mL heparin, 5 µM Y-27632, 1% L-Ala-Gln, and 1% P/S. For all cells that need to be preserved, use serum-free cell freezing medium (New Cell & Molecular Biotech, C40100) for storage.

### Lentiviral construction, packaging and infection

shRNA sequences were designed using BLOCK-iT™ RNAi Designer and cloned into the pLKO.1 vector (Addgene #8453). The vector was digested with AgeI (Abclonal, RK21125) and EcoRI (Abclonal, RK21102), ligated with annealed shRNA fragments using T4 DNA ligase (Abclonal, RK21501), and verified by Sanger sequencing. Knockdown efficiency was assessed in basal cells. For overexpression, *Efna3* CDS was cloned into pLeGO-SFFV-P2A-mCherry using the ClonExpress Ultra One-Step Cloning Kit (Vazyme, C115). Plasmids were prepared using an endotoxin-free plasmid extraction kit (Magen, P1112).

HEK293T cells were co-transfected with pMD.2, pspax2, and transfer plasmids using Eztrans (Shanghai Life-iLab Biotech Co., Ltd, AC04L099). Virus-containing medium was collected 60- and 88-hours post-transfection, concentrated by centrifugation at 27,000 rpm for 2 hours, and resuspended in DMEM/F12 with 10% FBS. Mammary organoids or cells were digested into single cells for infection. Lentivirus infection lasted 2 hours in ultra-low attachment culture dishes, and positive cells were sorted by flow cytometry. The shRNA sequences were as follows:

Scramble shRNA: 5’-CCTAAGGTTAAGTCGCCCTCG-3.’

*Efna3* shRNA: 5’-CCGGGGGTCTGCACTGTACATCTC-3’.

### Quantitative Real-Time PCR

RNA was extracted using an RNA extraction kit (Abclonal, RK30120) or Micro RNA extraction kit (Magen, R4125) and reverse transcribed into cDNA using ABScript III RT Master Mix for qPCR (Abclonal, RK20429). qPCR was performed with 2X Universal SYBR Green Fast qPCR Mix (Abclonal, RK21203) on a BIO-RAD CFX Connect Real-Time System. Actb was used as the internal reference. Primer sequences are listed in Supplementary Table 1.

### Immunofluorescence Staining

Cryosections of 6-week K14-GFP mouse were fixed in 4% paraformaldehyde (PFA) and permeabilized with 0.5% Triton X-100 in PBS for 35 minutes at room temperature. After permeabilization, samples were blocked in PBS containing 10% goat serum and 0.2% Tween-20 for 2 hours at room temperature or overnight at 4°C. Primary antibodies were incubated at 4°C overnight, followed by secondary antibody incubation for 2 hours at room temperature. Imaging was performed using a Nikon Spinning Disk Confocal Microscope. Primary antibodies used were ITGA2 antibody (HUABIO, #ET1611-57), ITGB1 antibody (R&D Systems, #MAB2405), ITGB3 antibody (HUABIO, #ET1606-49), and pFAK antibody (ThermoFisher, #44-624G).

### Colony Formation

A total of 500 luminal cells were cultured in ultra-low attachment 96-well plates(Jet Biofil) in stem cell medium: advanced DMEM/F12 containing 5% FBS, 10 ng/mL EGF, 20 ng/mL FGF2, 4 µg/mL heparin, 5 µM Y-27632, 1% L-Ala-Gln, 1% P/S, and 5% Matrigel. After 7 days of culture, colonies larger than 50 µm were counted. When FCCP (Selleck, S8276) was used, the final concentration was 0.5 nM.

### Mammary gland transplantation regeneration and visualization

For mammary gland regeneration transplantation, 3-week-old female nude mice were injected with 20,000 mammary epithelial cells into cleared fat pads. Mammary glands were collected after 4 weeks, tiled on adhesive glass slides, treated with Carnoy’s fixative solution, rehydrated, stained with carmine solution, dehydrated, and degreased before imaging. For breast tumor orthotopic transplantation, 6-week-old nude mice were injected with 100,000 tumor cells into the fat pad without removing the mammary gland tissue. Tumors from VO-PyMT cells were harvested after 3-4, and from PyMT cells after 8 weeks, ensuring that tumor sizes remained within animal welfare guidelines. Tumor mass was measured using a balance, and tumor volume was calculated with calipers using the formula: volume = length × width² × 0.5. Metastatic lungs were imaged using a Zeiss Axio Zoom V16 stereoscope.

### Image Acquisition and Time-Lapse Imaging

Confocal images were acquired using the Nikon CSU W1-01 SoRa+NIR Spinning Disk Microscope with CFI Plan Apochromat VC 20X or CFI Apo LWD 40XWI λS objectives. Images were processed with Fiji software. Time-lapse imaging was performed at 37°C with 5% CO₂, capturing images every 15 or 30 minutes. Basic medium as mentioned above was used for migration and live-cell imaging. DIC (differential interference contrast) images were acquired using the Zeiss observer z1 microscpoe with LD Plan-Neofluar 20x/0.4 Korr M27 objective. Time course imaging was performed at 24h,48h and 72h respectively. Images were processed with ImageJ software.

### Tracking of Organoids and Cells

ImageJ was used to measure organoid size, distance traveled, and speed, based on the center of gravity of the organoids. Green fluorescent-labeled cells were tracked automatically using the Spots function of Imaris software (Bitplane). The center of each cell was marked with a spot in each frame, and spots were connected over time. Mean cell speed was calculated by dividing the total track length by the tracking time, while displacement length represented the total cell displacement. An unpaired t-test was used to assess the statistical significance of cell speed and displacement lengths between the front and rear cells of the organoids.

### Cellular protrusion analysis

Protrusions were analyzed using 3D reconstructions of GFP-expressing organoids carrying the *R26R*^mT/mG^ reporter allele. Images were captured every 15 or 30 minutes over 10-15 hours and analyzed using Imaris (Bitplane). Protrusion data were collected by overlaying an eight-section pie chart with 45° intervals on a transparent circular grid. The 0 to 180° axis was aligned with the direction of organoid migration, and the pie center was placed over the center of each analyzed cell. The number of protrusions per bin per cell was counted every 15 or 30 minutes for a minimum of six hours. Protrusion data were plotted in sector charts using RStudio. A Chi-Squared test was used to determine the significance of a weighted mean direction, with the null hypothesis being that there is no mean direction.

### Bulk RNA Sequencing and Bioinformatics Analysis

For sequencing, basal (Ba), luminal (Lu), and basal-luminal (BaLu) cells were aggregated and subjected to a migration assay. After 55 hours of migration, cells were harvested using Recovery Solution (Corning, #354253) for subsequent RNA extraction. For the EFNA3 stimulation sequencing of luminal cells, luminal cells were serum-starved for 12 hours, followed by incubation with 1 µg/ml EFNA3 protein in serum-free medium for 24 hours prior to RNA extraction.

Total RNA was extracted using either the Micro RNA extraction kit (Magen, R4125) or the RNA extraction kit (Abclonal, RK30120) according to the manufacturers’ instructions. RNA quality was assessed using a 5300 Bioanalyzer (Agilent) and quantified with the ND-2000 spectrophotometer (NanoDrop Technologies). Only high-quality RNA samples (OD260/280 = 1.8–2.2, OD260/230 ≥ 2.0, RQN ≥ 6.5, 28S:18S ≥ 1.0, >10ng) were used for sequencing library construction.

RNA purification, reverse transcription, library construction, and sequencing were performed at Shanghai Majorbio Bio-pharm Biotechnology Co., Ltd. (Shanghai, China) according to the manufacturer’s protocols. RNA-seq transcriptome libraries were prepared using the Illumina® Stranded mRNA Prep, Ligation kit (San Diego, CA) with 30 ng of total RNA. Messenger RNA was isolated via poly-A selection using oligo(dT) beads and fragmented using a fragmentation buffer. Double-stranded cDNA was synthesized using the SuperScript double-stranded cDNA synthesis kit (Invitrogen, CA) with random hexamer primers. The cDNA was subjected to end-repair, phosphorylation, and adapter ligation according to library preparation protocols. Libraries were size selected for cDNA fragments of 300 bp using 2% Low Range Ultra Agarose, followed by PCR amplification with Phusion DNA polymerase (NEB) for 15 cycles. Libraries were quantified using a Qubit 4.0 fluorometer, and sequencing was performed on the NovaSeq 6000 platform with the NovaSeq Reagent Kit.

Raw paired-end reads were trimmed and quality-controlled using fastp with default parameters. Clean reads were aligned to the reference genome using HISAT2 software. Mapped reads were assembled for each sample using StringTie in a reference-based approach.

Raw reads in the fastq format were aligned to the mouse genome (mm10) using the subjunc function of the subread package (Version 2.0.3). Gene counts for each sample were generated from GTF annotation files (gencode.vM30.annotation.gtf) using the FeatureCounts function of the subread package. Differential gene expression was analyzed using the DESeq2 R package, and gene set enrichment analysis (GSEA) was conducted with the clusterProfiler R package. Ligand-receptor interaction data were obtained from (github.com/arc85/celltalker) using the celltalker R package.

### Bioinformatics Analysis of Genes in Different Stages of Mammary Gland Development

scRNA-seq data covering different stages of mammary gland development were obtained from GSE164017. Data input, quality control, integration, and cell clustering were performed using the Seurat R package. Basal cells were marked by Krt14 and Krt5 expression, while luminal cells were identified by Krt18 and Krt8 expression. Gene expression levels were averaged across different developmental stages in both basal and luminal cells.

scRNA-seq data containing TEB, duct-derived basal, and luminal cells for cell junction’s pathway and gene expression analysis are derived from GSE164017. Differential gene analysis was performed using DESeq2 and GO enrichment analysis and GSEA were conducted with the clusterProfiler R package.

Data on EFNA3 mRNA expression in breast cancer subtypes were based on PRJNA762594 and its correlation with oxidative phosphorylation as well as gene expression were based on The Cancer Genome Atlas (TCGA) Breast Cancer (BRCA) datasets, available at UCLAN (http://ualcan.path.uab.edu/analysis.html). The distal metastasis-free survival curves (DMFC) are based on 5287 breast cancer patients. Kaplan-Meier Plotter (http://kmplot.com/analysis/index.php?p = service&cancer = breast) was used for the analysis.

### Statistical Analysis

Sample sizes are indicated in the figure legends. Statistical significance between groups was assessed using two-tailed Student’s t-tests. For sector charts, a two-sample Chi-squared test was used. Error bars represent either standard deviation (SD) or standard error of the mean (SEM), with statistical significance denoted as follows: *P < 0.05, **P < 0.01, ***P < 0.001, ****P < 0.0001. Non-significant results are indicated as n.s.

### Data Availability

All data supporting the conclusions of the paper are available in the article and corresponding figures. scRNA-seq data and human breast cancer data were downloaded from public databases as indicated in the methods section.

## ACKNOWLEDGEMENTS

We thank members of the Lu lab for discussions and Drs. Chenleng Cai, and Yikang Rong for thoughtful comments and suggestions. We thank technical support from the Molecular Imaging Core Facility, and Molecular and Cell Biology Core Facility in Hengyang Medical School at South China University. We especially thank Ms. Rui Wang from the Molecular Imaging Core Facility in School of Life Science and Technology at ShanghaiTech University for her advice on image processing. This work was supported by grants from the National Science Foundation of China (32470886 to P.L., and 82330035 to K.X.).

## AUTHOR CONTRIBUTIONS

J.C. Data acquisition, analysis, and curation

R.M. Data acquisition, Data quality control, analysis, diagrams, and review.

D.F. and Z.D. Data analysis, bioinformatics analysis.

1. Y. Lu, O.D.K., and K.X., intellectual contributions, review.

P.L., Conceptualization, funding acquisition, data curation, and writing.

