## Supplementary Figure for "Basal EFNA3 Facilitates Luminal-Driving Mammary Epithelial Migration and Cancer Metastasis by Promoting OXPHOS"

**Supplementary Fig. 1. Basal Cells Exhibit Lower Adhesion and Higher Speed than Luminal Cells During Collective Migration**

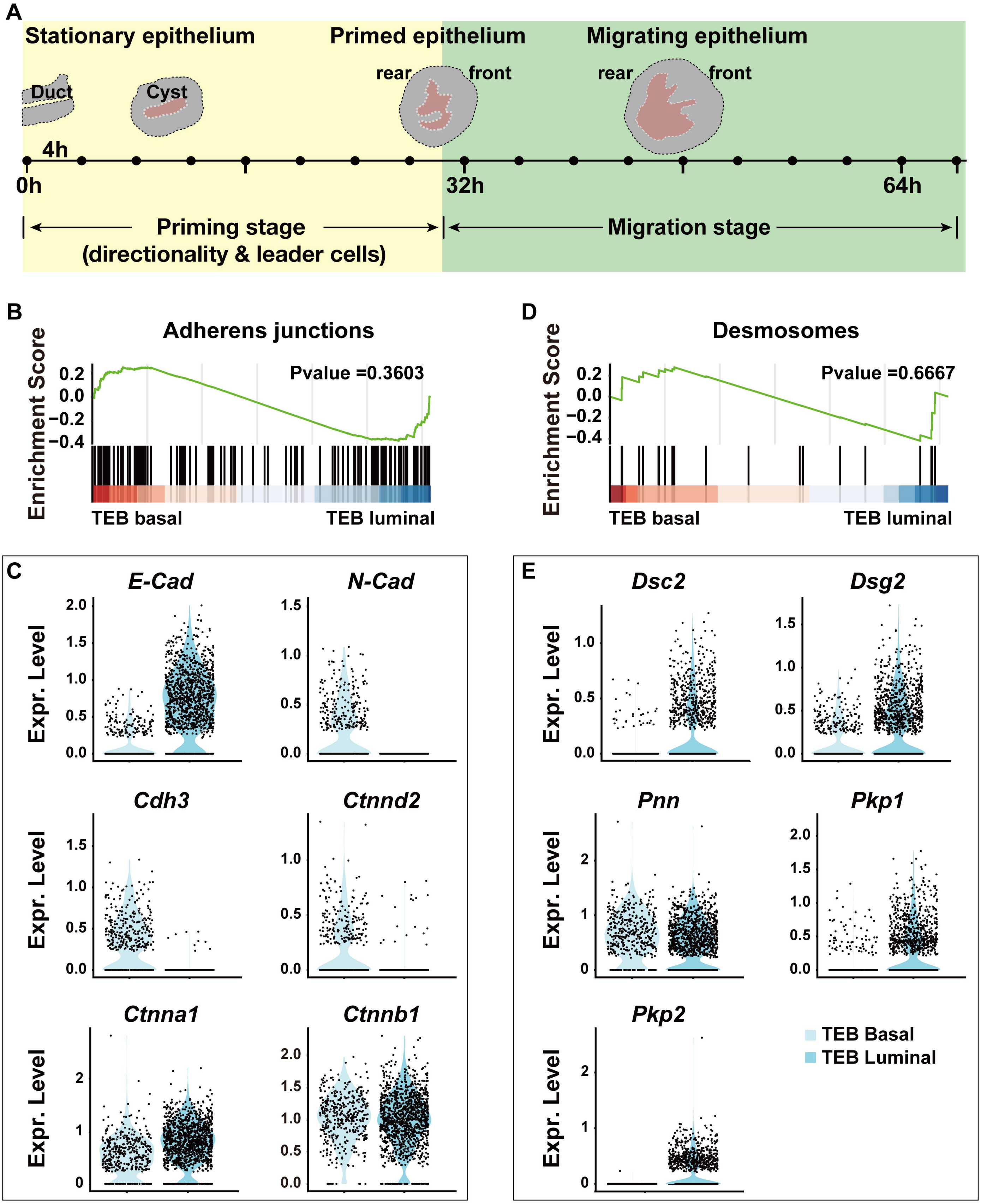

**Supplementary Fig. 2. The focal adhesion pathway is active in luminal cells.**

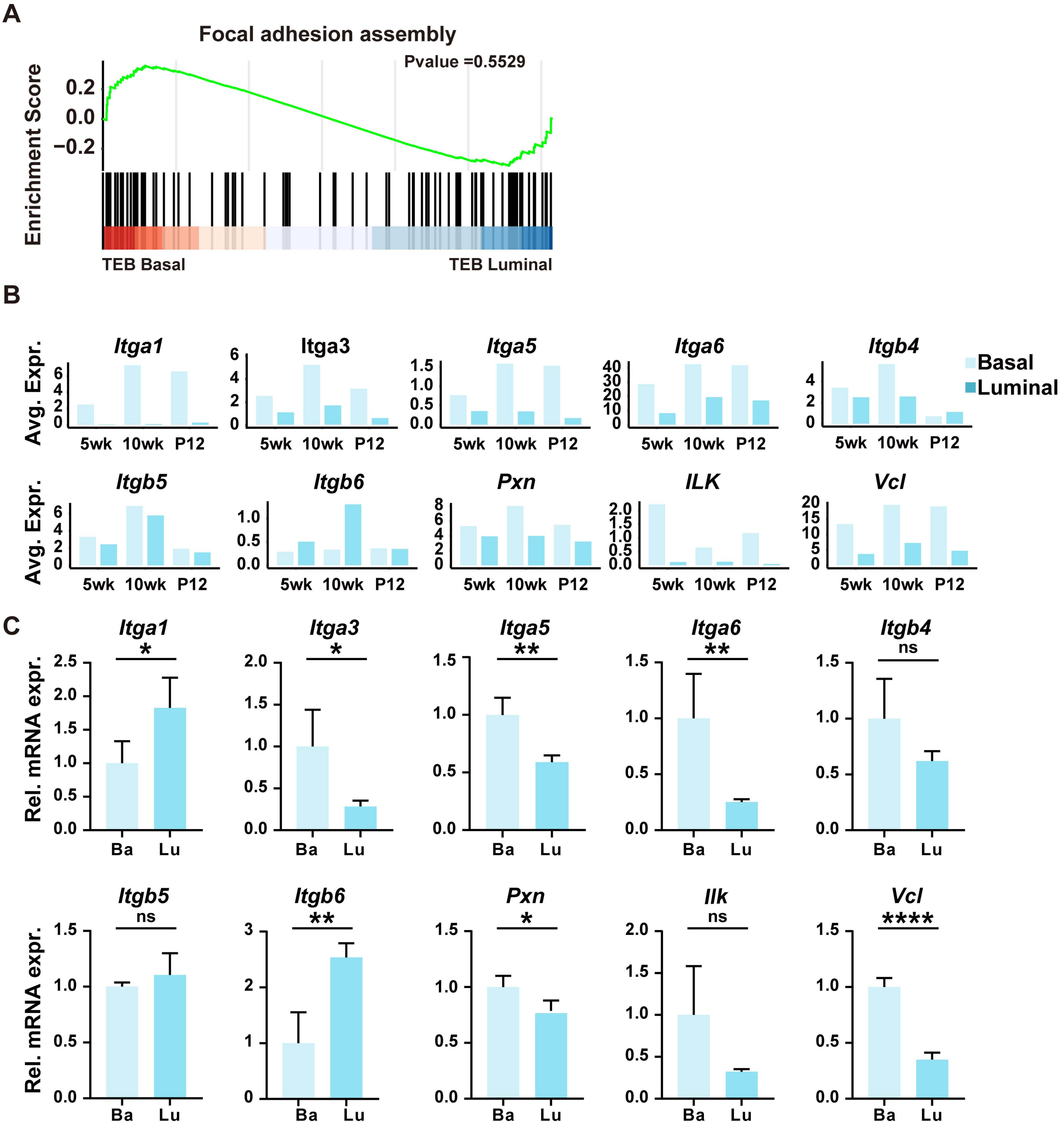

**Supplementary Fig. 3. EFNA3 receptor expression.**

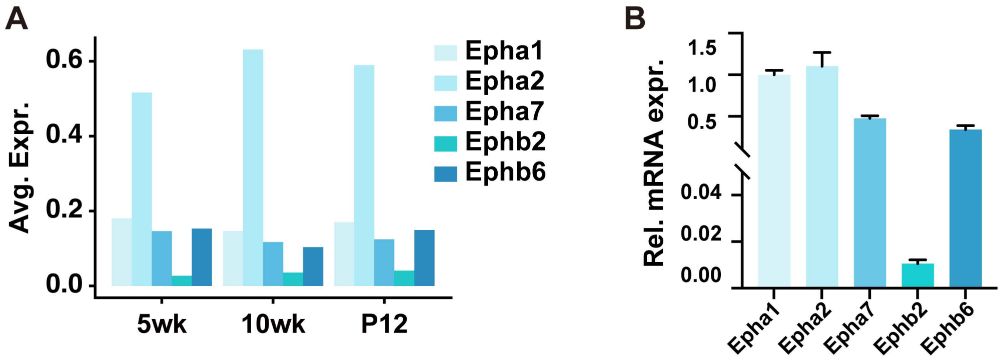

**Sup.Fig. 4. Basal EFNA3 Promotes Luminal Migration In Vitro and Epithelial Branching In Vivo by Enhancing OXPHOS.**

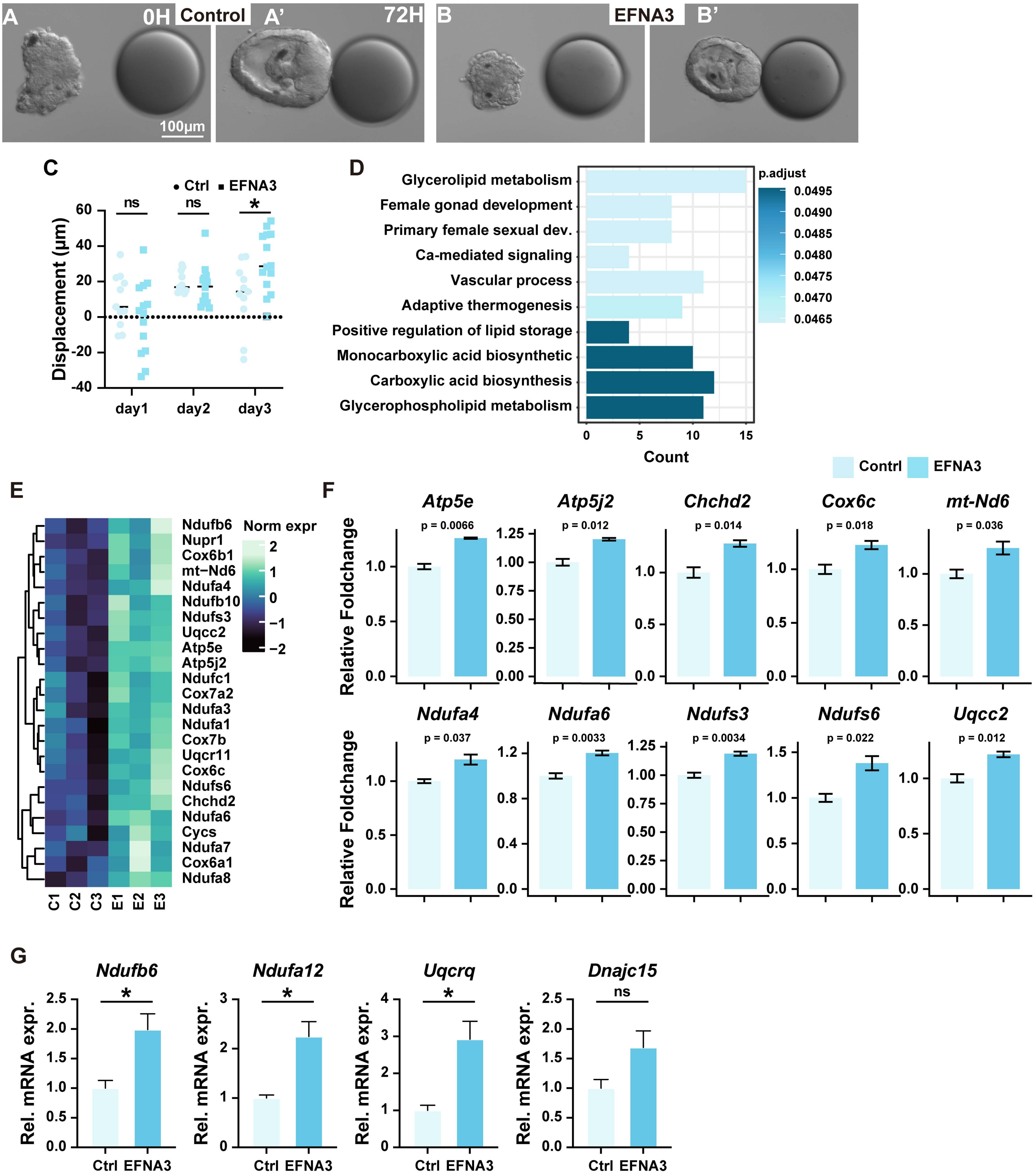

**Supplementary Fig. 5. EFNA3 as a Prognostic and Therapeutic Target to Inhibit Breast Cancer Metastasis.**

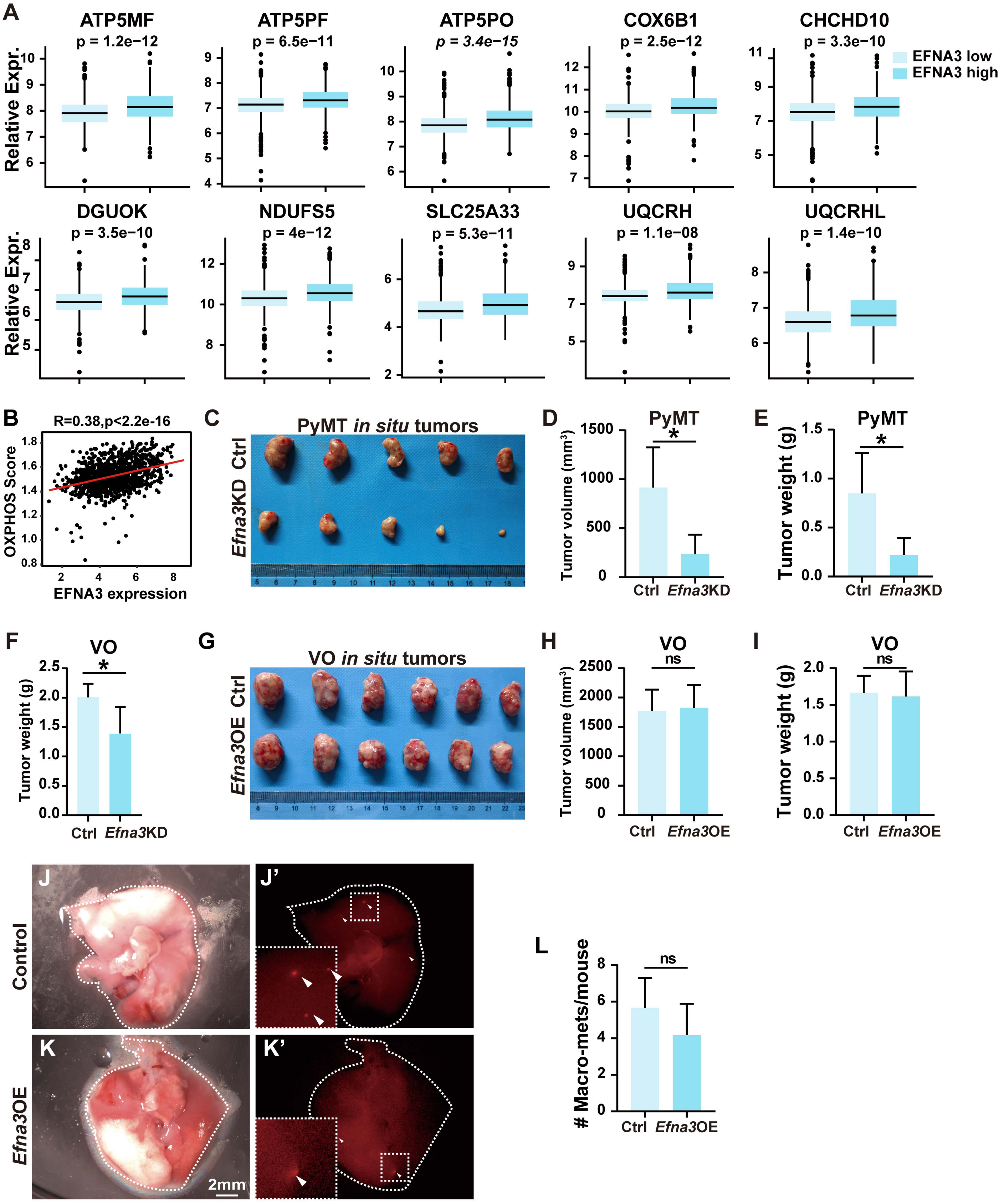
