## Supplementary Figure Legends for "Basal EFNA3 Facilitates Luminal-Driving Mammary Epithelial Migration and Cancer Metastasis by Promoting OXPHOS"

**Supplementary Figure 1: Basal Cells Exhibit Lower Adhesion and Higher Speed than Luminal Cells During Collective Migration.**

(**A**) Schematic of two stages of epithelial migration based on our recent findings[5]. First, an FGF10 gradient promotes preferential cell proliferation at the front of the stationary organoid epithelium, establishing "front-rear" polarity. As a result, the organoid becomes primed, with the front region stratified and lacking apicobasal polarity, leading to the generation of frontal leader cells. In the second stage, the primed organoid epithelium becomes migratory and moves toward the FGF10 signal, with leader cells driving the process.

(**B**-**E**) GSEA analyses and expression profiles of individual cells within the adherens junction (**B**, **C**) and desmosome (**D**, **E**) pathways in basal and luminal cells of TEBs. Note that the P values in (**B**) and (**D**) are not significant.

Abbreviations: h, hour; Expr, expression.

**Supplementary Figure 2: The Focal Adhesion Pathway Is Active in Luminal Cells.**

(**A**) GSEA analysis of the focal adhesion pathway comparing basal and luminal cells in TEBs. Note that the P values were not significant.

(**B**) mRNA expression of selected genes in the focal adhesion pathway at various stages of mouse mammary gland development, based on scRNA-seq dataset analysis (GSE164017).

(**C**) Quantitative PCR (qPCR) analysis showing mRNA expression levels of selected genes in the integrin signaling pathway in basal and luminal cells of the branching mammary gland at 8 weeks of development.

Data are presented as mean $\pm$ SD (with over 500 cells counted per sample). n.s., not significant, $P \geqslant 0.05$; $*P < 0.05$; $**P < 0.01$; $****P < 0.0001$.

Abbreviations: Avg, average; Rel, relative; Expr, expression; Ba, basal; Lu, luminal; wk, week; P, pregnancy day.

**Supplementary Figure 3: EFNA3 receptor expression.**

(**A**) EFNA3 receptor mRNA expression levels in luminal cells across different stages of mouse mammary gland development, based on analysis of scRNA-seq datasets (GSE164017).

(**B**) Quantification of EFNA3 receptor mRNA expression levels in luminal cells from branching mammary glands at the 8-week stage, as determined by qPCR.

Abbreviations: Avg, average; Expr, expression; wk, week; P, pregnancy day.

**Supplementary Figure 4: EFNA3 promotes OXPHOS.**

(**A**-**C**) The effect of EFNA3 protein on FGF10-induced epithelial organoid migration. EFNA3 was added to the medium at concentrations of 0 μg/ml (**A**, **A'**; control) or 1 μg/ml (**B**, **B'**; experimental). Migration effects were quantified (**C**).

(**D**) Gene Ontology analysis of the main pathways associated with downregulated genes in luminal cells following EFNA3 stimulation.

(**E**) Heat map showing upregulation of oxidative phosphorylation-related genes after EFNA3 stimulation.

(**F**) mRNA expression levels of selected genes in the OXPHOS pathway based on RNA sequencing data.

(**G**) Quantification of OXPHOS pathway gene expression using qPCR in luminal cells treated with EFNA3 at concentrations of 0 μg/ml (control) or 1 μg/ml (experimental).

Data are presented as mean ± SD. n.s., not significant, $P \geqslant 0.05$; $*P < 0.05$.

Abbreviations: dev, development; C1-3, control groups 1-3; E1-3, EFNA3 groups 1-3; Rel, relative; expr, expression.

**Supplementary Figure 5: EFNA3 as a Prognostic and Therapeutic Target to Inhibit Breast Cancer Metastasis.**

(**A**) mRNA expression levels of selected genes in the OXPHOS pathway in breast cancer tissues with either high or low EFNA3 expression.

(**B**) Positive correlation between EFNA3 expression and OXPHOS activity. Higher EFNA3 expression is associated with increased oxidative phosphorylation.

(**C**-**E**) Tumor growth from PyMT cells 8 weeks post-surgery in nude mice with reduced *Efna3* expression via shRNA knockdown (**C**), with tumor volume (**D**) and weight (**E**) quantified.

(**F**) Quantification of tumor weight from VO-PyMT cells 4weeks post-surgery in nude mice, comparing control to *Efna3* knockdown.

(**G**-**I**) Tumor growth from VO-PyMT cells 3 weeks post-surgery in nude mice with either control or *Efna3* overexpression (**G**), with tumor volume and weight quantified in (**H**) and (**I**), respectively.

(**J**-**L**) Lung metastasis of VO-PyMT cells transfected with lentivirus expressing control (**J**, **J'**) or *Efna3*OE (**K**, **K'**), visualized in wholemount lung tissue under brightfield (**J**, **K**) or fluorescence stereoscope (**J'**, **K'**). (**L**) Quantification of lung metastasis from VO-PyMT cells transplanted into the cleared fat pad 3weeks post-surgery.

Data are presented as mean $\pm$ SD. n.s. = not significant ($P \geqslant 0.05$); $*P < 0.05$.

Abbreviations: Expr, expression; macro-mets, macro-metastases.
